# Characterisation of growth plate dynamics in murine models of osteoarthritis

**DOI:** 10.1101/2020.10.14.339119

**Authors:** Hasmik J. Samvelyan, Kamel Madi, Anna E. Törnqvist, Behzad Javaheri, Katherine A. Staines

**Author notes:** Corresponding author Hasmik Jasmine Samvelyan, University of Brighton, Lewes Road, Brighton BN2 4GJ.

## Abstract

**Background:** The purpose of this study was to investigate growth plate dynamics in surgical and loading murine models of osteoarthritis, to understand whether abnormalities in these dynamics predict osteoarthritis vulnerability.

**Methods:** 8-week-old C57BL/6 male mice underwent destabilisation of medial meniscus (DMM) (*n =* 8) surgery in right knee joints. Contralateral left knee joints had no intervention (controls). In 16-week-old C57BL/6 male mice (*n* = 6), osteoarthritis was induced using non-invasive mechanical loading of right knee joints with peak force of 11N. Non-loaded left knee joints were internal controls. Chondrocyte transiency in tibial articular cartilage and growth plate was confirmed by histology and immunohistochemistry. Tibial subchondral bone parameters were measured using microCT and correlated to 3D GP bridging analysis.

**Results:** Higher expression of chondrocyte hypertrophy markers; Col10a1 and MMP13 were observed in tibial articular cartilage chondrocytes of DMM and loaded mice. In tibial growth plate, Col10a1 and MMP13 expressions were widely dispersed in a significantly enlarged zones of proliferative and hypertrophic chondrocytes in DMM (*p*=0.002 and *p*<0.0001, respectively) and loaded (both *p*<0.0001) tibiae of mice compared to their controls. 3D quantification revealed enriched growth plate bridging and higher bridge densities in medial compared to lateral tibiae of DMM and loaded knee joints of the mice. Growth plate dynamics were associated with higher subchondral bone volume fraction (BV/TV; %) in medial tibiae of DMM and loaded knee joints and epiphyseal trabecular bone volume fraction in medial tibiae of loaded knee joints.

**Conclusions:** The results confirm articular cartilage chondrocyte transiency in a surgical and loaded murine model of osteoarthritis. Herein, we reveal for the first time spatial variation of growth plate bridging in surgical and loaded osteoarthritis models and how these may contribute to anatomical variation in vulnerability of osteoarthritis development.

## Background

Osteoarthritis (OA) is a chronic musculoskeletal disease and a leading cause of disability and major healthcare costs in the world. It is estimated that worldwide 10% of men and 18% of women aged over 60 years have symptomatic OA (1) and, therefore, the predicted increase in the ageing population and longevity will result in a greater occurrence of the disease. OA is a complex disease in which the pathogenesis, cellular and molecular mechanisms of initiation and progression are not completely understood. It is characterised by progressive loss of articular cartilage (AC), formation of osteophytes, subchondral bone (SCB) sclerosis, synovial proliferation, inflammation and lax tendons. These can ultimately lead to a loss of joint function, pain, reduced mobility and disability (2). Despite the significant healthcare and economic burden there are few non-invasive therapies available to patients. Therefore, understanding the pathogenesis of OA and defining the molecular mechanisms underpinning AC degeneration can lead to the development of successful targeted and effective disease-modifying treatments.

Primary OA is described as naturally occurring OA affecting one joint (localised) or three or more joints (generalised), while secondary OA is associated with various causes and risk factors leading to the disease including trauma, obesity, diabetes, metabolic bone and congenital disorders (3). AC degeneration is one of the main hallmarks of OA and previous research has largely sought to identify mechanisms underpinning its deterioration. Fully developed, uncalcified AC is populated by a single resident cell chondrocytes, which maintain a stable phenotype characterised by small cell size and expression of tenascin-C (4). The inherent stability of AC chondrocytes ensures that dynamic events are restricted to assure lifelong articular integrity and healthy joint function. In contrast, epiphyseal growth plate (GP) chondrocytes have a transient phenotype to ensure long bone development (endochondral ossification) and growth. GP chondrocytes undergo a differentiation sequence of proliferation, maturation and hypertrophy. The final stage of chondrocyte hypertrophy enables mineralisation of the cartilage extracellular matrix, vascular invasion and subsequent replacement of the mineralised cartilage anlagen with bone (5). These processes are coupled, however, with sexual maturation the human GP undergoes progressive narrowing as bony bridges form and span its width. This ultimately leads to complete GP closure and cessation of human growth. Indeed, in humans the longitudinal bone growth stops with the onset of puberty, the metaphysis then fuses with the epiphysis and the GP disappears (6). In mice, longitudinal bone growth does not cease at sexual maturity instead it slows dramatically at puberty, but the GPs do not completely fuse and disappear (7).

We have previously shown that in the STR/Ort mouse, a naturally occurring OA murine model, AC chondrocytes transform from their inherently stable phenotype to a transient one, characteristic of the chondrocytes in the GP. This was confirmed by immunolabelling for chondrocyte hypertrophy markers; type X collagen (Col10a1) and matrix metalloproteinase 13 (MMP13)(8). Further, we revealed accelerated longitudinal bone growth, aberrant expression of GP markers (Col10a1 and MMP13) and increased GP chondrocyte maturation in these mice. Consistent with this, using a novel synchrotron computed tomography method we revealed enriched GP bone bridging in STR/Ort mouse tibiae indicative of advanced GP closure which may underpin OA (8,9).

Despite this, interlinks between the differing chondrocyte phenotypes in the AC and the GP, and the precise contribution that the GP and its fusion mechanisms may play in underpinning OA vulnerability is not yet understood. The aim of this study was to investigate whether surgically (destabilisation the medial meniscus [DMM]) or non-surgically (mechanical loading) induced OA onset in C57BL/6 mice is linked to altered GP dynamics. Indeed, understanding this will inform strategies for maintaining musculoskeletal health in ageing by potentially identifying whether GP dynamics may predict who is at risk of OA in later life and ultimately developing targeted OA treatments.

## Methods

### Animals

Male C57BL/6 wild type mice at 7 weeks of age (young adult) were obtained from Charles River Laboratories Inc. (Margate, UK). The mice were acclimatised to their surroundings for seven days. All mice were allowed free access to water and maintenance diet ad libitum (Special Diet Services, Witham, UK) in a 12-hour light/dark cycle at a room temperature of 21 ± 2^0^C and relative humidity of 55 ± 10%.

### The destabilisation of the medial meniscus (DMM)

8-week old wild type C57/BL6 male mice underwent DMM surgeries to induce OA-like changes in the right knee joints under isoflurane-induced anaesthesia (*n* = 8/group). We chose not to performed SHAM (placebo) surgery on the left contralateral knee of the animals based upon animal welfare grounds in keeping with the 3Rs, since it was previously shown that there is no difference in OA scores between SHAM-operated and non-operated knee joints (10,11). Animals were monitored for unexpected adverse effects to reduce suffering and distress. Following transection of the medial meniscotibial ligament (MMTL) to destabilise the medial meniscus, the skin was closed and anaesthesia reversed (10). Eight weeks later, mice were culled by exsanguination and confirmation of death by cervical dislocation. The knee joints of all the mice were dissected, fixed in 4% paraformaldehyde for 24 hours at 4°C, and then stored in 70% ethanol.

### In vivo loading of the knee joint

The right knee joints of 16-week-old wild type C57BL/6 male mice (*n* = 6) were subjected to non-invasive, dynamic axial mechanical loading under the isofluorane-induced anaesthesia (liquid isofluorane was vaporised to a concentration of 4% and maintained at a concentration of 2% with oxygen) for 7 min/day, 3 alternate days a week for 2 weeks according to the protocols described in the previous studies (12,13). The left knee joints were non-loaded internal controls in these animals. Briefly, using a servo-electric materials testing machine (Electroforce 3100, Bose, UK), axial compressive loads were applied through the right knee joint via customised concave cups which held the knee and ankle joints flexed and the tibiae vertically.

The tibia was held in place by continuous static preload of 0.5N onto which dynamic loads were superimposed in a series of 40 trapezoidal shaped waveform cycles with steep up and down ramps and a peak force of 11N for 0.05 seconds (0.025 seconds rise and fall time; 9.9 seconds baseline hold time between periods of peak loading). The right and left knees were dissected 3 days after the final loading episode. Mice were culled by exsanguination and confirmation of death by cervical dislocation. Knee joints were fixed in 4% paraformaldehyde for 24 hours at 4°C before being stored in 70% ethanol.

### Micro-computed (microCT) tomography and 3-dimensional (3D) bridging analysis

Scans were performed with an 1172 X-Ray microtomograph (Bruker MicroCT, Kontich, Belgium) to evaluate the SCB and GP bridging. High-resolution scans with an isotropic voxel size of 5 µm were acquired (50 kV, 200 µA, 0.5 mm aluminium filter, 0.6° rotation angle). The projection images were reconstructed and binarised with a threshold of 0 to 0.16, ring artefact reduction was set at 10 and beam hardening correction at 0% using the SkyScan NRecon software package (v1.6.9.4, Bruker MicroCT). The images then were realigned vertically using DataViewer software (v1.5.1.2 64-bit, Bruker MicroCT) to ensure similar orientation for analysis. Hand-drawn regions of interests (ROI) of the SCB and epiphyseal trabecular bone for each tibial lateral and medial compartments were selected (14). The structural parameters of tibial SCB plate and epiphyseal trabecular bone were calculated using 3D algorithms of SkyScan CTAn software (Bruker MicroCT) including SCB (SCB BV/TV; %) and trabecular bone volume fraction (Tb. BV/TV; %) and correlated to GP bridging analysis using a 3D synchrotron-computed tomography quantification method as previously described (15). Briefly, microCT scans of the tibiae were segmented using a region-growing algorithm within the Avizo® (V8.0, VSG, Burlington, VT, USA) software. The central points of each bony bridges were identified and projected on the tibial joint surface. The distribution of the areal number density of bridges (N, the number of bridges per 256 µm × 256 µm window; *d* = *m*/*V*) is then calculated and superimposed on the tibial joint surface (each bridge has a colour that represents the areal number density at the bridge location). The SCB plate and epiphyseal trabecular bone thickness (Th; mm) was determined and colour-coded thickness images were generated using the Avizo® software(15).

### Histological analysis

The left and right knee joints of all the mice were decalcified in 10% ethylenediaminetetraacetic acid (EDTA) solution, wax-embedded at Leica EG1160 Tissue Embedding Station and 6 μm coronal sections cut using Leica RM2135 manual microtome.

### Immunohistochemistry

Immunohistochemical analysis of chondrocyte transiency markers in tibial AC and GP was performed on 6 μm coronal sections from the middle region of the knee joint, using anti-matrix metalloproteinase 13 (anti–MMP13) (1:200 dilution; Abcam) or anti-collagen type X (anti-Col10a1) (1:100 dilution; Abcam) antibodies. As a control, an equal concentration of rabbit IgG was used. For immunohistochemical localisation of MMP13 and Col10a1, sections were dewaxed in xylene and rehydrated. Sections were incubated at 37 °C for 30 min in 1mg/ml trypsin for antigen demasking. Endogenous peroxidases were blocked by treatment with 0.3% H_2_O_2_ in methanol (Sigma) for 30 min at room temperature. The Vectastain ABC universal detection kit (Vector Laboratories, Peterborough, UK) was used to detect the biotinylated secondary antibody (Anti-Mouse IgG Reagent) after incubation for 30 min at room temperature according to the manufacturer’s instructions. Diaminobenzidine (DAB) solution used to detect the location of antigens. The sections were finally dehydrated, counterstained with haematoxylin and mounted in DePeX. All sections to be compared were immunolabelled at the same time to standardise conditions and minimise any differences in antibody incubation times. For each group, immunolabelling was performed on four individual animals per experimental group and representative images taken using a light microscope.

### Growth plate (GP) zone analysis

The width of the GP proliferating and hypertrophic zones, as well as the total GP width, were measured at 10 different points along the length of the GP in 6μm coronal sections from the middle region of the knee joint of four individual animals per experimental group, using a light microscope and ImageJ software.

### Statistical analysis

All analyses were performed with GraphPad Prism software 6.0f version (GraphPad Inc, La Jolla, CA, USA) using a two-sided 0.05 level of significance. All analyses were conducted blindly to minimise the effects of subjective bias. The results were presented as the mean ± standard error of the mean (SEM). The Normal distribution of data was assessed using the Shapiro-Wilk normality test. For comparing two groups (experimental with control, or medial with the lateral compartment of tibiae), two-tail Student’s *t*-test (paired or unpaired) was used. For comparing more than two groups, two-way ANOVA (analysis of variance) was used with Tukey post-hoc test.

## Results

### Transient chondrocyte behaviour in the tibial AC of DMM and loaded C57BL/6 young adult male mice

We first sought to confirm whether loss of AC in C57BL/6 mice with surgically and loading induced OA was associated with the expression of markers of transient chondrocyte phenotype. In accordance with previous studies, immunohistochemistry analysis showed higher expression levels of well-established chondrocyte hypertrophy markers; Col10a1 and MMP13, observed in tibial AC of C57BL/6 mice that have undergone DMM surgery or mechanical loading compared with non-operated and non-loaded control left tibiae, respectively (Fig. 1a and b). The expression pattern of Col10a1 was largely restricted to hypertrophic chondrocytes in the uncalcified zone of the AC of unaffected condyles of non-operated (Fig. 1a) and non-loaded (Fig. 1b) mouse left joints as expected (16). Whereas, the immunolabeling of Col10a1 was more widespread throughout the extracellular matrix (ECM) of the AC in affected right joints of DMM (Fig. 1a) and loaded (Fig. 1b) mice. Similarly, immunohistochemistry analysis showed positive MMP13 labelling in both superficial and deep articular chondrocytes in the right joints of DMM and loaded C57BL/6 male mice compared to the control knee joints (Fig. 1a and b). These findings confirm an aberrant deployment of transient chondrocytes in uncalcified AC.

**Fig. 1.**
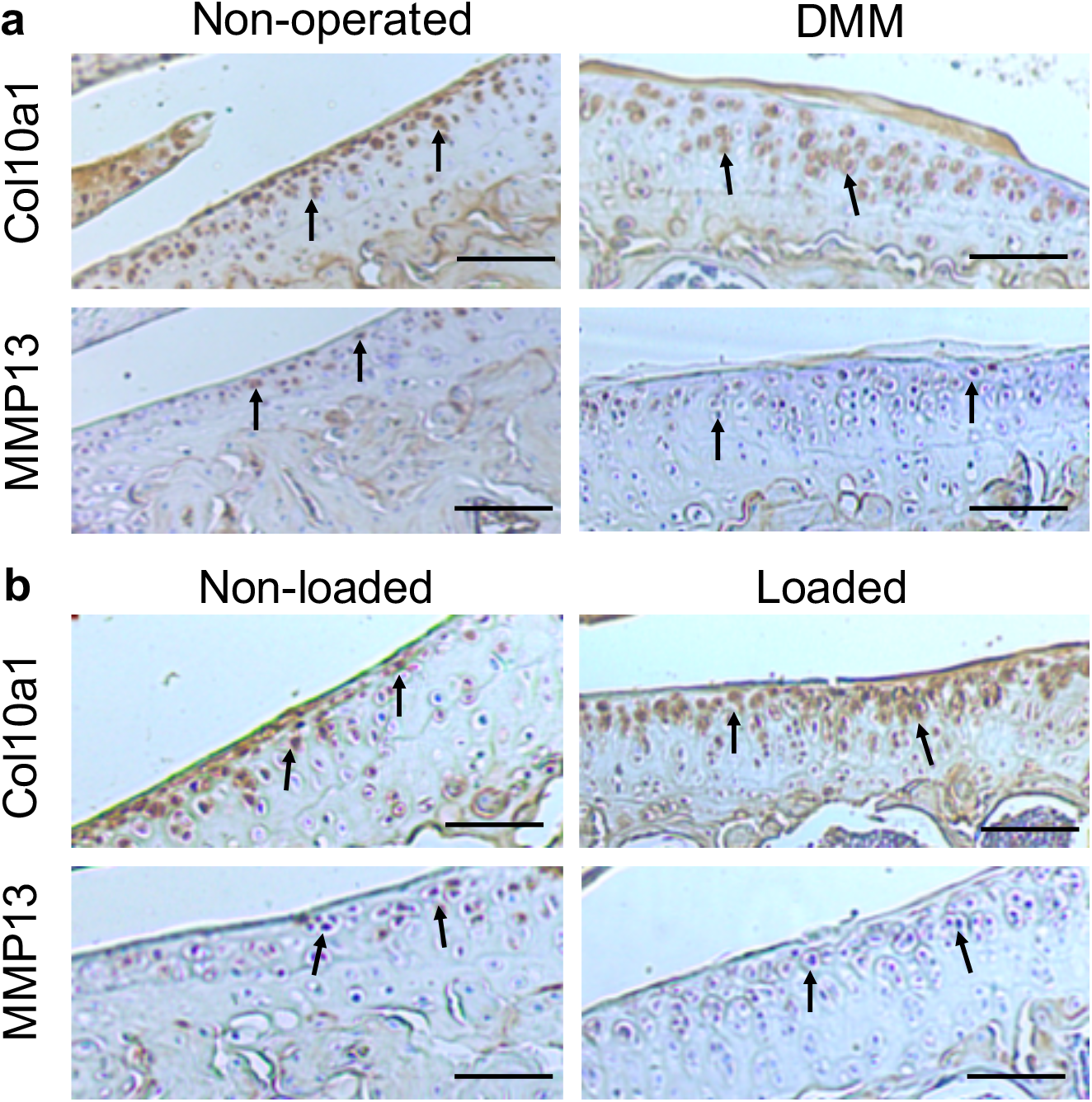
Immunohistochemical labelling in the tibial articular cartilage of mouse knee joints. Immunohistochemical labelling for type X collagen (Col10a1) and matrix metalloproteinase (MMP13) in the articular cartilage of non-operated and DMM (**a**), or non-loaded and mechanically loaded (**b**) knee joints of C57BL/6 male mice. Arrows indicate examples of positive staining in the tibiae. Images are representative of results in 3 individual mice. Scale bar = 200μm

### Dysfunctional GP dynamics in DMM and loaded tibiae of C57BL/6 young adult male mice

To determine the GP phenotype in these OA models, we first completed histological assessment of GP zones. Our analysis revealed significantly enlarged proliferative and hypertrophic zones of chondrocytes in both DMM (*p*=0.002 and *p*<0.0001, respectively) and loaded (both *p*<0.0001) tibiae of C57BL/6 mice compared to their controls, and significantly increased total GP width (both *p*<0.0001) (Fig. 2a). This is suggestive of aberrant GP dynamics in these OA models.

**Fig. 2.**
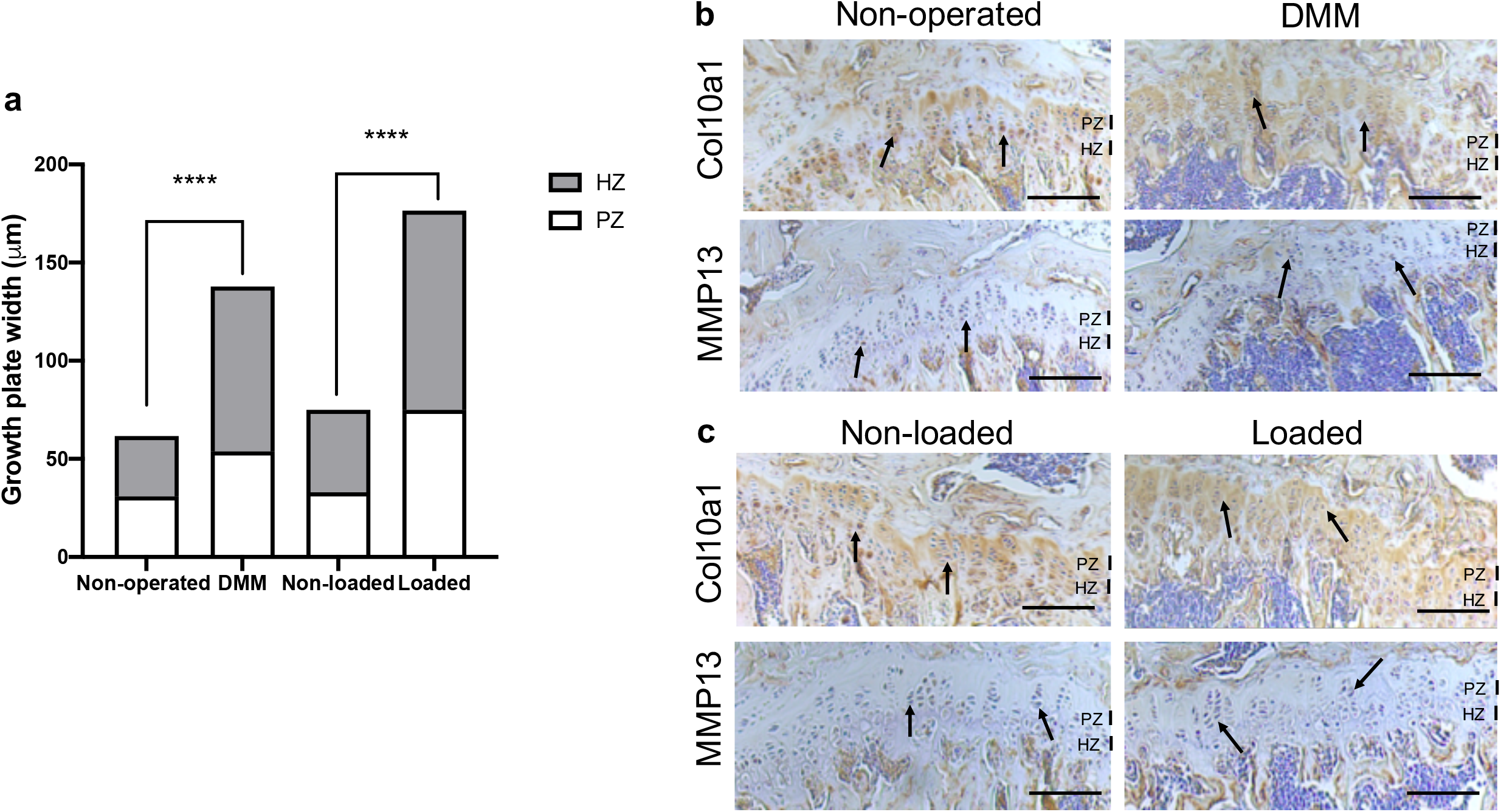
Growth plate dynamics in the tibial growth plate of mouse knee joints. Growth plate zone width of non-operated and DMM, or non-loaded and loaded knee joints of C57BL/6 male mice (**a**). Ten measurements per section were obtained along the length of the tibial growth plate in the middle region of the knee joint (n=4 mice for each experimental group). Immunohistochemical labelling for type X collagen (Col10a1) and matrix metalloproteinase (MMP13) in the growth plate of DMM and non-operated (**b**), or mechanically loaded and non-loaded (**c**) knee joints of C57BL/6 male mice. Arrows indicate examples of positive staining in the tibiae. Images are representative of results in 4 individual mice. Scale bar = 200μm. PZ = proliferative zone; HZ = hypertrophic zone **** *p*<0.0001

We next determined the expression of known chondrocyte transiency markers in the GPs. In the tibial GP of mice with surgically and loading induced OA, Col10a1 expression was more greatly and widely dispersed throughout the zones of proliferative and hypertrophic chondrocytes compared with their controls (Fig. 2b and c). Indeed, immunolabeling for Col10a1 revealed the expected localisation in the GP of non-operated and non-loaded mouse tibiae, limited primarily to the hypertrophic zone and underlying adjacent metaphyseal bone (Fig. 2b and c). This disrupted the distribution of a GP zone marker was also evident for MMP13 in the GP of DMM and loaded C57BL/6 mouse tibiae compared to their controls (Fig. 2b and c). Together, the results may indicate associations between dysfunctional GP morphology and marker expression, and disease development in these murine models of OA.

### Associations between GP bridging in DMM and loaded tibiae of C57BL/6 young adult male mice and OA development

To further correlate aberrant longitudinal GP dynamics, GP bridging and OA development in these models, we used our newly developed 3D method to quantify GP bone bridges across the tibial epiphysis of DMM and loaded mice for the first time (Fig. 3). 3D quantification revealed a significantly higher number of GP bridges in medial compared to lateral tibiae that underwent DMM surgeries (196 ± 33 versus 306 ± 32; *p*=0.03) (Fig. 3b and e). This significant difference was not observed in non-operated tibiae (254 ± 32 versus 326 ± 32; *p*=0.14) (Fig. 3a and e). Similarly, significantly enriched GP bridging was evident in the medial compartment of loaded tibiae in comparison to the lateral compartment (539 ± 36 versus 745 ± 18; *p*=0.0009) (Fig. 3d and e), and in those of non-loaded tibiae, although less pronounced than in the loaded right knee joints (596 ± 22 versus 726 ± 44; *p*=0.04) (Fig. 3c and e). However, no significant differences in GP bridge numbers and densities were observed between interventions (DMM versus non-operated, and loaded versus non-loaded) at this time point in either the medial or lateral compartment. These results were consistent with the areal bridge density analysis. The mean areal bridge densities were significantly greater in medial compared to the lateral compartment of DMM (8.2 ± 0.9 versus 10.3 ± 0.9; *p*=0.01) and loaded tibiae (16.3 ± 0.6 versus 20.6 ± 0.5; *p*<0.0001) (Fig. 3f). However, no significant differences in the mean areal bridge densities were observed between interventions (DMM versus non-operated, and loaded versus non-loaded).

**Fig. 3.**
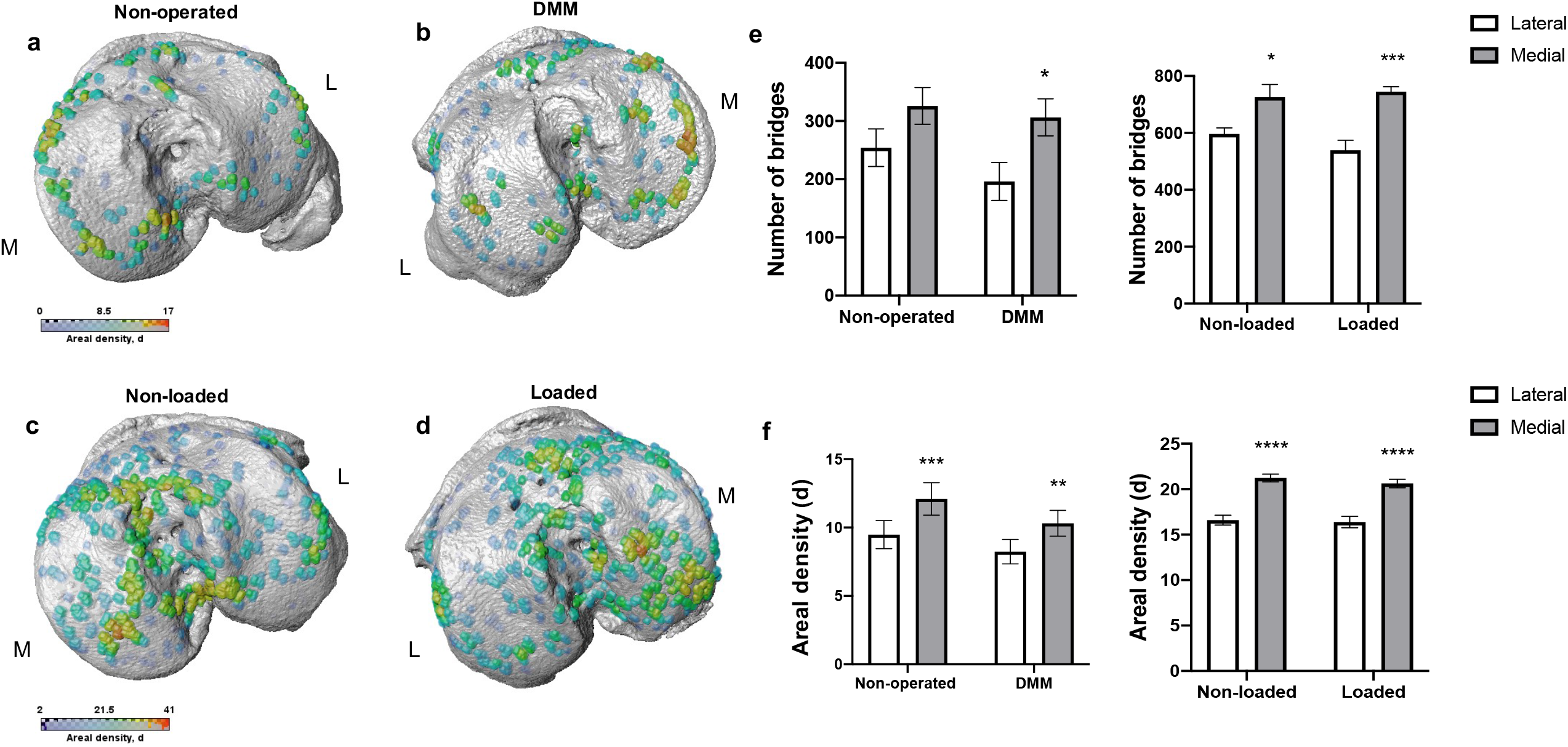
Bridge location and bridge areal densities across the tibial growth plate of mice. Location and areal densities of bridges across the growth plate projected on the medial (M) and lateral (L) tibial joint surface in non-operated (**a**), DMM (**b**), non-loaded (**c**) and loaded (**d**) tibiae of mice at 16 and 18 weeks of age, number of bridges in lateral and medial tibiae of non-operated compared to DMM and non-loaded compared to loaded tibiae of mice (**e**), Areal density (d) of bridges in medial compared to lateral tibiae, defined as the number of bridges per 256 mm x 256 mm window of non-operated and DMM or non-loaded and loaded knee joints (**f**). Bars represent mean ± SEM. Group sizes were *n* = 8 for non-operated and DMM-operated mice and *n* = 6 for non-loaded and loaded mice. * *p*<0.05 ** *p*<0.01 *** *p*<0.001 **** *p*<0.0001

Anatomical variations were observed however in all samples, with focal high density clusters forming in the anterior region of the medial loaded tibiae versus those more in the central and posterior of non-loaded tibiae (Fig. 3c and d). Similarly, in the DMM tibiae, GP bridges were more widespread across the tibiae than in non-operated which were predominantly observed around the periphery (Fig. 3a and b). Together, these results suggest that murine OA models exhibit spatial variations in GP bridging in comparison to controls.

### MicroCT analysis and 3D visualisation of SCB plate and epiphyseal trabecular bone

To establish whether spatial variations in GP bridging are associated with the SCB plate and epiphyseal trabecular bone abnormalities after the DMM surgery and mechanical loading in our mice, we performed microCT analysis and determined local thickness of SCB plate and epiphyseal trabecular bone. The SCB plate volume fraction (SCB BV/TV) was significantly higher in medial compared to lateral compartment of DMM and non-operated tibiae (DMM: SCB BV/TV 29.4 ± 3.1% versus 36.7 ± 4.5%, *p*=0.03, non-operated: SCB BV/TV 35.7 ± 0.7% versus 42.6 ± 1.5%, *p*=0.004) (Fig. 4a). No significant differences were observed between DMM and non-operated tibia (Fig. 4a), or in the epiphyseal trabecular bone volume fraction (Tb. BV/TV; Fig. 4b). Conversely, in the loaded and non-loaded tibiae, epiphyseal trabecular bone volume fraction (Tb. BV/TV) was significantly increased in the medial compared to lateral compartment (loaded: Tb. BV/TV 60.7 ± 1.8% versus 73.4 ± 2.8%, *p*=0.005, non-loaded: Tb. BV/TV 61.4 ± 2.2% versus 77.4 ± 2.5%, *p*=0.0005). No significant differences were observed between interventions (Fig. 4d). The SCB plate volume fraction (SCB BV/TV) was significantly higher in medial compared to lateral compartment of loaded tibiae (40.1 ± 1.7% versus 47.1 ± 1.6%, *p*=0.02), but not that of non-loaded tibiae (42.1 ± 1.1% versus 45.8 ± 1.6%, *p*=0.34) or between interventions (Fig. 4c).

**Fig. 4.**
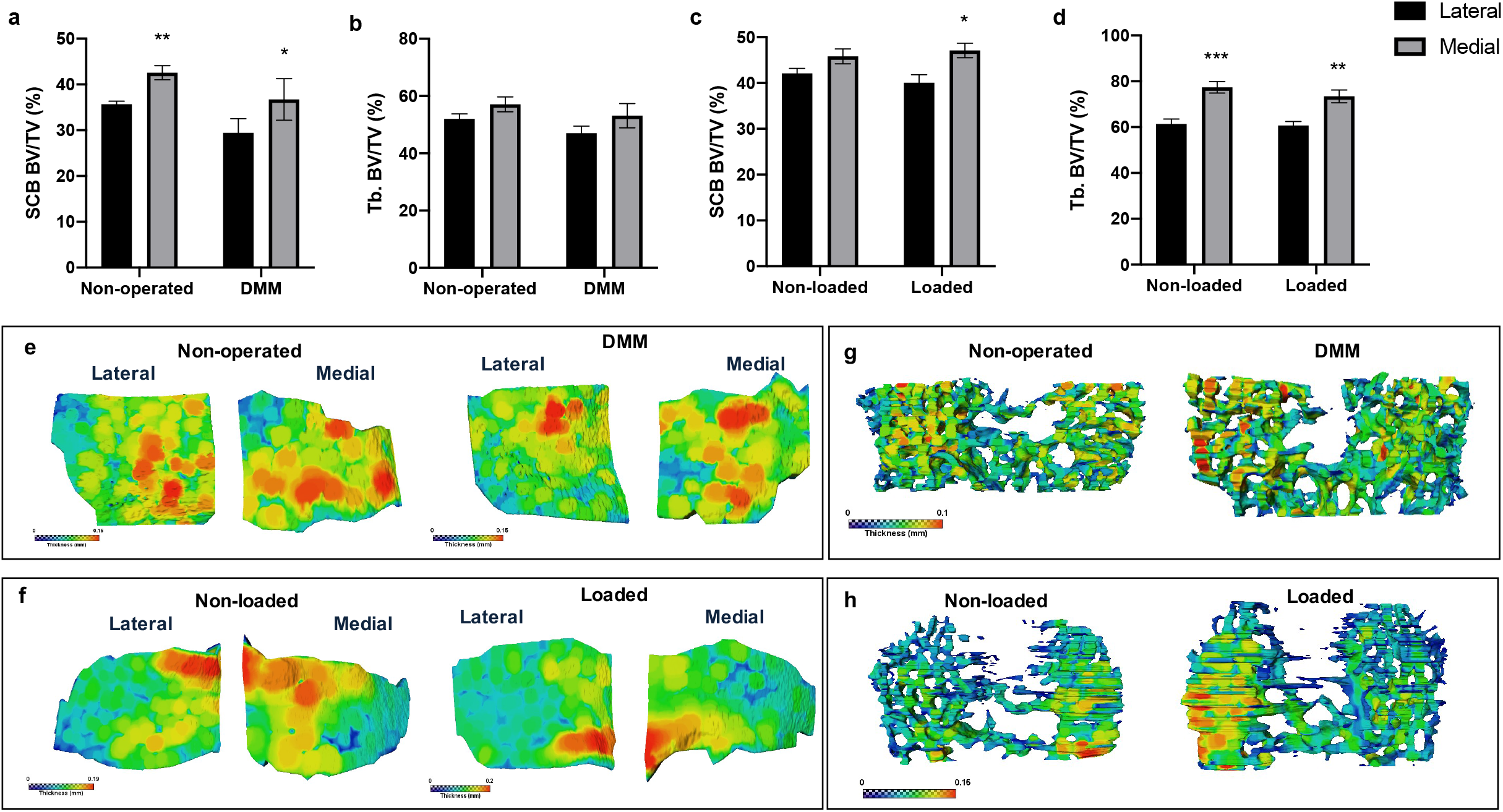
MicroCT analysis of epiphyseal region of medial and lateral tibiae of mice. MicroCT analysis of the epiphyseal region of the lateral and medial tibiae in non-operated controls and DMM-operated knee joints subchondral bone volume fraction (SCB BV/TV) (**a**) and epiphyseal trabecular bone volume fraction (Tb. BV/TV) (**b**). MicroCT analysis of the epiphyseal region of the medial and lateral tibiae in non-loaded controls and loaded knee joints subchondral bone volume fraction (SCB BV/TV) (**c**) and epiphyseal trabecular bone volume fraction (Tb. BV/TV) (**d**). Representative colour coded images of lateral and medial subchondral bone plate thickness of non-operated and DMM-operated tibiae of mice (**e**). Representative colour coded images of lateral and medial subchondral bone plate thickness of non-loaded and loaded tibiae of mice (**f**). Representative colour coded images of epiphyseal trabecular bone thickness of non-operated and DMM-operated (**g**), or non-loaded and loaded tibiae of mice (**h**). Bars represent mean ± SEM. Group sizes were *n* = 8 for non-operated and DMM-operated mice and *n* = 6 for non-loaded and loaded mice. * *p*<0.05 ** *p*<0.01 *** *p*<0.001

Colour-coded SCB plate thickness analysis revealed anatomical variation in SCB plate thickness between DMM and non-operated, and loaded and non-loaded plates (Fig. 4e and 4f). Differences in the non-loaded and loaded SCB plates were particularly apparent (Fig. 4f), and correlated with the clusters of higher density of GP bridges previously observed (Fig. 3c and d). Epiphyseal trabecular bone thickness was not significantly altered in response to either invasive or non-invasive intervention (data not shown; Fig. 4g and h).

## Discussion

Together, the data presented herein reveals altered GP dynamics in both surgical (DMM) and non-invasive loading murine *in vivo* models of OA. Our data confirm previous studies showing changes in AC of the knee joints of these mice consistent with the aberrant deployment of chondrocyte transiency, and reveal disrupted GP morphology, increased GP chondrocyte differentiation indicated by widespread expression of chondrocyte hypertrophy markers and increased GP zone widths in these models. Moreover, we have utilised a recently developed 3D model to discover for the first time anatomical variations in GP bone bridging, indicative of GP fusion, which are spatially correlated with increase in SCB thickness. These data reveal that altered GP dynamics and spatial differences in GP bridging may contribute to an anatomical variation in vulnerability to OA development in surgical and loaded murine models of OA.

which with findings described above may indicate an endochondral defect in AC and GP cartilage in these mouse models of OA. These findings build upon our previous work in the STR/ort OA murine model which is predisposed to developing spontaneous idiopathic OA whilst its nearest available parental strain, the CBA mouse, has a very low susceptibility which makes them effective controls for the studies (17). We have previously shown that aberrant deployment of transient chondrocyte behaviour, consistent with re-initiation of endochondral processes, occurs in uncalcified AC of STR/ort mouse knee joints compared to CBA controls (8). Here we extend these studies to look at the expression of transient markers in other stratifications of OA using murine models. Surgically induced DMM model is widely used for target validation studies or evaluation of the pathophysiological roles of many molecules in OA. Following DMM, medial displacement of the medial meniscus in a mouse knee joint provides a smaller area to transmit the weight-bearing forces and leads to an increased local mechanical stress (18). Whereas, in a cyclic AC tibial compression model, the non-invasive dynamic mechanical loading applied to the mouse tibia through the knee and ankle joints, modifies AC structure locally through a mechanoadaptive homeostatic response contributing to OA development (19). Hypertrophic chondrocytes in the calcified cartilage and GP of the healthy joints express Col10a1 (17,19). The calcified cartilage acts to protect the uncalcified AC through maintaining its ECM in an unmineralised state and the stability of the AC. However, hypertrophic differentiation of these chondrocytes contributes to AC matrix degradation, calcification and vascular invasion resulting in the demise of the AC (23). Consistent with this, the expression of Col10a1, as examined using immunohistochemistry, has been observed throughout the AC in the joints of our both DMM and loaded mice. Further, the higher expression level of another marker of chondrocyte hypertrophy; MMP13 has been detected in superficial and uncalcified chondrocytes in the AC of DMM and loaded mice compared to their non-operated and non-loaded left knee joints. Indeed, cartilage degradation observed in OA has been attributed to an elevated production of proteolytic enzymes among which MMP13 has a major role (24,25). Studies using transgenic mice deficient in catabolic transcription factors that induce hypertrophic differentiation revealed that animals were protected against surgically and chemically induced OA further highlighting the role of transient chondrocytes in AC degradation (26,27).

The results of the present study indicate that in these murine OA models, there is a significantly enlarged zone of GP proliferative and hypertrophic chondrocytes, compared with the chondrocyte zones in the GP of control tibiae of both models. Similarly, a significantly increased cumulative GP width was observed in these models. This is associated with aberrant widespread expression of Col10a1 and MMP13 in the GP of DMM and loaded tibiae of C56BL/6 mice, compared to the GP of non-operated and non-loaded control knees. Longitudinal bone growth is determined by the modifying number of chondrocytes in the proliferative zone of GP, rate of their proliferation, the extent of chondrocyte hypertrophy and controlled synthesis and degradation of ECM throughout the GP (28). Altered growth rate and mechanical modulation of GP function appear to result from complex interactions of changes in the states of these chondrocytes, as does the rate of GP closure due to the formation of bone bridges forming and spanning the width of the GP.

Our report of aberrant GP chondrocyte dynamics in these DMM and loaded C57BL/6 mice is further strengthened by our data acquired using a 3D quantification method of bone bridging across the tibial epiphysis (15). Here we show significantly enriched spatial localisation of GP bone bridging clustering and number in the tibial medial compared to the lateral compartments of DMM mice in comparison non-operated mice. We reveal focal clusters of higher density bridges in different anatomical regions of the tibiae, which was correlated with increased SCB thickness. This is consistent with our previous work in the STR/Ort mouse model of spontaneous OA in which we observed similar spatial variations. We postulate that the formation of these bridges may be accelerated by local factors like instigating altered mechanoadaptive response and that these spatial variations in GP bridging may disclose the anatomical vulnerability to OA. These findings are supported by the previous studies suggesting associations between local mechanical stress caused by medial displacement of the medial meniscus (29) or cyclic tibial AC compression (19) and GP function in C57BL/6 mice. They are further supported by our previous work in which we revealed, by finite element modelling, that GP bridges act to dissipate stresses upon loading to the overlying SCB and thus suggest that this contributes to OA seen in these models (15). Indeed the GP is subject to a variety of mechanical forces placed upon it, and strain distributions are inhomogeneous through the GP with the hypertrophic zone exhibiting low mechanical stiffness. The precise interplay between GP bridging and mechanics has yet to be determines, specifically in the context of OA and how this may increase OA vulnerability.

Identification of modified GP dynamics underpinning human OA would allow elucidation into OA predisposition and ultimately enable the development of novel and specific therapeutic interventions. Our recent work in the MRC National Survey of Health and Development (NSHD) found that increased height in childhood was associated, albeit modestly, with lower odds of knee OA at age 53 years, as was adult achieved height (30). Concurrent with this, a recent study from offspring in the Avon Longitudinal Study of Parents and Children (ALSPAC) found height tempo (corresponding to pubertal timing) to be strongly associated with the hip shape models which may be related to hip future risk OA(31). Further, several known OA susceptibility single nucleotide polymorphisms (SNPs) have been associated with hip shape in perimenopausal women in the ALSPAC(32). Most intriguingly, when combined with data from other cohorts, the eight SNPs independently associated with hip shape were intimately associated with the process of endochondral ossification (33). Together these data provide further tantalising evidence that there are associations between GP dynamics during adolescence and OA development.

The limitation of this study is that we only examined the GP at one specific time point for each model and therefore more time points are required to fully understand how these GP bridges temporally affect SCB changes and OA pathology. With its controllability, the intermittent non-invasive mechanical loading model will allow in future to distinguish between short- and long-term effects of various cyclic loading regimens on SCB and trabecular bone parameters as well as AC integrity, GP dynamics and allow correlation of these to initiation and progression of human OA. Indeed, it is known that in the loading model, the short-term intervention is not sufficient to induce significant changes in subchondral thickness and thus longer intervention times for both the DMM and loading models may also prove useful in pursuit of understanding these relationships (14).

## Conclusions

Our studies indicate that aberrant GP dynamics may contribute to OA pathology in both surgical and loading mouse OA models, and that similar GP-related osteoarthritic pathological changes happen in different *in vivo* models of secondary (post-traumatic) OA (18) through inducing direct (DMM) or indirect (mechanical loading) injuries to the joints. We reveal for the first time spatial variation of GP bridging in these OA models which may contribute to anatomical variation in vulnerability of OA development. Our GP bony bridging analysis may signify accelerated cartilage-bone transition in these affected joints, advancing our understanding of GP closure mechanisms and how these contribute to the health of the joint. Further, our work yields more insights into the changes in the micro-mechanical environment of the GP and chondrocytes within the GP. This work extends our knowledge on the contribution of the GP in OA development, which is vital if we are to reduce the burden of this global disease.

## List of abbreviations

AC: articular cartilage
Col10a1: type X collagen
DAB: Diaminobenzidine
DMM: destabilisation of medial meniscus
ECM: extracellular matrix
GP: growth plate
MMP13: matrix metalloproteinase 13
MMTL: meniscotibial ligament
OA: osteoarthritis
SCB: subchondral bone

## Declarations

### Ethics approval and consent to participate

All procedures complied with the United Kingdom Animals (Scientific Procedures) Act 1986 and were approved by The University of Edinburgh Roslin Institute’s Animal Users and Research Ethics Committees.

### Consent for publication

Not applicable

### Availability of data and materials

The datasets used and/or analysed during the current study are available from the corresponding author on reasonable request.

### Authors’ contributions

All authors contributed to the study design, analysis and interpretation of the data, drafted, critically reviewed, edited and approved the version of the manuscript for publication.

## Acknowledgements

The authors would like to thank The Roslin Institute of The University of Edinburgh for the assistance with the care of animal models in this study.

